# Cis acting variation is common, can propagates across multiple regulatory layers, but is often buffered in developmental programs

**DOI:** 10.1101/2020.05.21.107961

**Authors:** Swann Floc’hlay, Emily Wong, Bingqing Zhao, Rebecca R. Viales, Morgane Thomas-Chollier, Denis Thieffry, David A. Garfield, Eileen EM Furlong

**Affiliations:** Institut de Biologie de l’ENS (IBENS), École Normale Supérieure, CNRS, INSERM, Université PSL, 75005 Paris, France; Molecular, Structural and Computational Biology Division, Victor Chang Cardiac Research Institute, Darlinghurst, New South Wales, Australia; School of Biotechnology and Biomolecular Sciences, UNSW Sydney, Kensington, New South Wales, Australia; European Molecular Biology Laboratory (EMBL), Genome Biology Unit, D-69117, Heidelberg, Germany; Institut Universitaire de France (IUF), 75005 Paris, France

**Keywords:** Natural sequence variation, developmental enhancers, F1 embryos, open chromatin, transcription, gene regulation, selection

## Abstract

Precise patterns of gene expression are driven by interactions between transcription factors, regulatory DNA sequence, and chromatin. How DNA mutations affecting any one of these regulatory ‘layers’ is buffered or propagated to gene expression remains unclear. To address this, we quantified allele-specific changes in chromatin accessibility, histone modifications, and gene expression in F1 embryos generated from eight *Drosophila* crosses, at three embryonic stages, yielding a comprehensive dataset of 240 samples spanning multiple regulatory layers. Genetic variation in *cis*-regulatory elements is common, highly heritable, and surprisingly consistent in its effects across embryonic stages. Much of this variation does not propagate to gene expression. When it does, it acts through H3K4me3 or alternatively through chromatin accessibility and H3K27ac. The magnitude and evolutionary impact of mutations is influenced by a genes’ regulatory complexity (*i*.*e*. enhancer number), with transcription factors being most robust to *cis*-acting, and most influenced by *trans*-acting, variation. Overall, the impact of genetic variation on regulatory phenotypes appears context-dependent even within the constraints of embryogenesis.

## Introduction

The development of a multicellular organism requires the tight regulation of gene expression in both space and time to ensure that reproducible phenotypes are obtained across individuals and environmental conditions. DNA regulatory elements (*e*.*g*. promoters and enhancers) are essential to this process by integrating regulatory information from sequence-specific transcription factors (TFs), RNA polymerase II (Pol II), and other regulatory proteins to drive specific spatio-temporal patterns of expression during development. But while gene expression patterns are typically quite precise, the DNA regulatory elements that control such expression states are replete with genetic variation (mutations) that can impact transcriptional regulation at multiple levels including TF binding (Kasowski et al. 2010; Spivakov et al. 2012; Behera et al. 2018), chromatin state (Waszak et al. 2015), transcriptional start site usage (Schor et al. 2017), gene expression levels (Garfield et al. 2013; Battle et al. 2015), and transcript isoform diversity (Cannavo et al. 2017). As a result, genetic variation in regulatory elements can contribute to variation in disease susceptibility among individuals (Epstein 2009; Lowe and Reddy 2015) and to evolutionary change between species (Wittkopp and Kalay 2011), by impacting higher-level phenotypes.

Although regulatory mutations can have large effects, many behave effectively neutrally, making it challenging to predict which genetic variants will have an impact. Part of the difficulty comes from the general lack of knowledge about which regions of non-coding DNA have regulatory (not just biochemical) function. An additional challenge is the apparent robustness of gene regulatory networks. At least within a laboratory context, whole sections of regulatory DNA can be removed with little apparent impact on phenotype or fitness (Ahituv et al. 2007), and evolutionarily divergent regulatory sequences are often swapped between species with few detectable changes in gene expression (Borok et al. 2010). These studies demonstrate that developmental systems have the ability to compensate or “buffer” the effects of regulatory mutations, *e*.*g*. via compensation by other regulatory elements with partially overlapping activities (Hong et al. 2008; Frankel et al. 2010; Cannavo et al. 2016).

The complex relationship between DNA sequences and regulatory function further complicates our understanding of how mutations can impact gene regulation. For example, mutations affecting TF binding motifs can have a large impact on chromatin accessibility, Pol II occupancy, histone modifications and gene expression (Kircher et al. 2019). But in some contexts/tissues, TF binding is driven by collective processes that can include protein-protein as well as protein-DNA interactions, such that mutations affecting a single TF motif may not substantially affect TF recruitment (Junion et al. 2012; Doitsidou et al. 2013; Uhl et al. 2016; Khoueiry et al. 2017). Moreover, many sequence variants affecting TF occupancy *in vivo* lie outside the TF’s binding motif, and are likely due to variation affecting the binding of co-occurring factors (Kasowski et al. 2010; Zheng et al. 2010; Reddy et al. 2012) or an overall change in DNA shape (Lu and Rogan 2018). To make matters more complex, enhancer output is not a strict function of all TF’s occupancy – enhancers often contain binding sites for multiple factors with redundant input, and in some cases, different combinations of TFs can produce the same expression output (Brown et al. 2007; Zinzen et al. 2009; Khoueiry et al. 2017). Even in cases where an enhancer’s activity is abolished by mutations, the gene’s expression may not be affected, as genes often have many enhancers with partially overlapping activity, which can buffer the functional impact of genetic variation impacting a single enhancer (Hong et al. 2008; Frankel et al. 2010; Cannavo et al. 2016). With a few exceptions (Bullaughey 2011), this complex genotype-to-phenotype relationship cannot be modelled using regulatory sequence information alone, but rather must be evaluated empirically (Khoueiry et al. 2017).

Allelic-specific data provides a unique opportunity to study the molecular mechanisms of *cis*-acting variation and has uncovered multiple regulatory processes through which cis-acting variation impacts transcriptional control (Kilpinen et al. 2013; Chen et al. 2016). F1 crosses of inbred strains provide an elegant method to determine the contribution of both *cis* and *trans* variation by overcoming the limits of genetic variation between trios and the general lack of statistical power to interrogate *trans*-acting variation using population data (Wittkopp et al. 2004; Tirosh et al. 2009; Goncalves et al. 2012; Wong et al. 2017). Taking advantage of this F1 design, we set out to better understand how natural sequence variation impacts gene regulation during embryonic development. We collected *Drosophila* F1 hybrid embryos and quantified allele-specific changes in open chromatin (ATAC-Seq), enhancer and promoter activity (using H3K27ac or H3K4me3 & H3K27ac ChIP-Seq as proxies, respectively), and gene expression (RNA-seq). Our half-sibling design of F1 embryos was generated by crossing males from eight genetically distinct, wild-derived isogenic lines from the *Drosophila* Genetic Reference Panel (DGRP) (Mackay et al. 2012) to females from a common, laboratory-derived isogenic reference strain. In addition to having practical advantages for conducting large scale crosses, as described below, the use of a common female line allowed us to evaluate the impact of regulatory mutations while controlling for maternal effects, which can contribute disproportionately to variability in early developmental phenotypes (Lynch and Walsh 1998; Garfield et al. 2013). By collecting matched phenotypic measurements from two parental strains (*F0*), we also estimated the heritability of *cis*-acting mutations and the relative magnitude of *trans*-acting genetic variation that contributes to phenotypic divergence.

Overall, we find allelic variation in chromatin accessibility and histone marks to be common and significantly correlated between regulatory layers, with the effects of regulatory mutations being more strongly coupled at promoters than distal elements (putative enhancers). Using this genetic variation as a perturbation to gene regulation, we uncovered different mechanistic rules in the order of information flow during transcriptional regulation. Specific classes of genes, such as TFs, are in general more strongly buffered against the effects of this variation, which in turn reflects their patterns of inheritance and genetic architecture (having a greater proportion of *trans* and less additive heritability). In some cases, selection is driven to near fixation in gene expression (but interestingly not in upstream regulatory layers), affecting genes involved in environmental responses and pesticide resistance. Taken together, this comprehensive data set provides new insights into the functional impact of *cis*-regulatory DNA variation and how this is transmitted across different regulatory layers during embryogenesis, and how patterns of inheritance can influence the visibility of regulatory sequence variants to natural selection.

## Results

### Quantifying gene expression and regulatory element activity in hybrid embryos

We collected F1 hybrid embryos by mating eight genetically distinct inbred lines from the DGRP collection (Mackay et al. 2012) to females from a common maternal line (Fig. 1a). The resulting F1 panel contains an average of 567,412 SNPs per cross, and a total of 1,455,988 unique SNPs covering a range of minor allele-frequencies and levels of conservation (phyloP scores) (Fig. S1a, Table S1).

**Figure 1:**
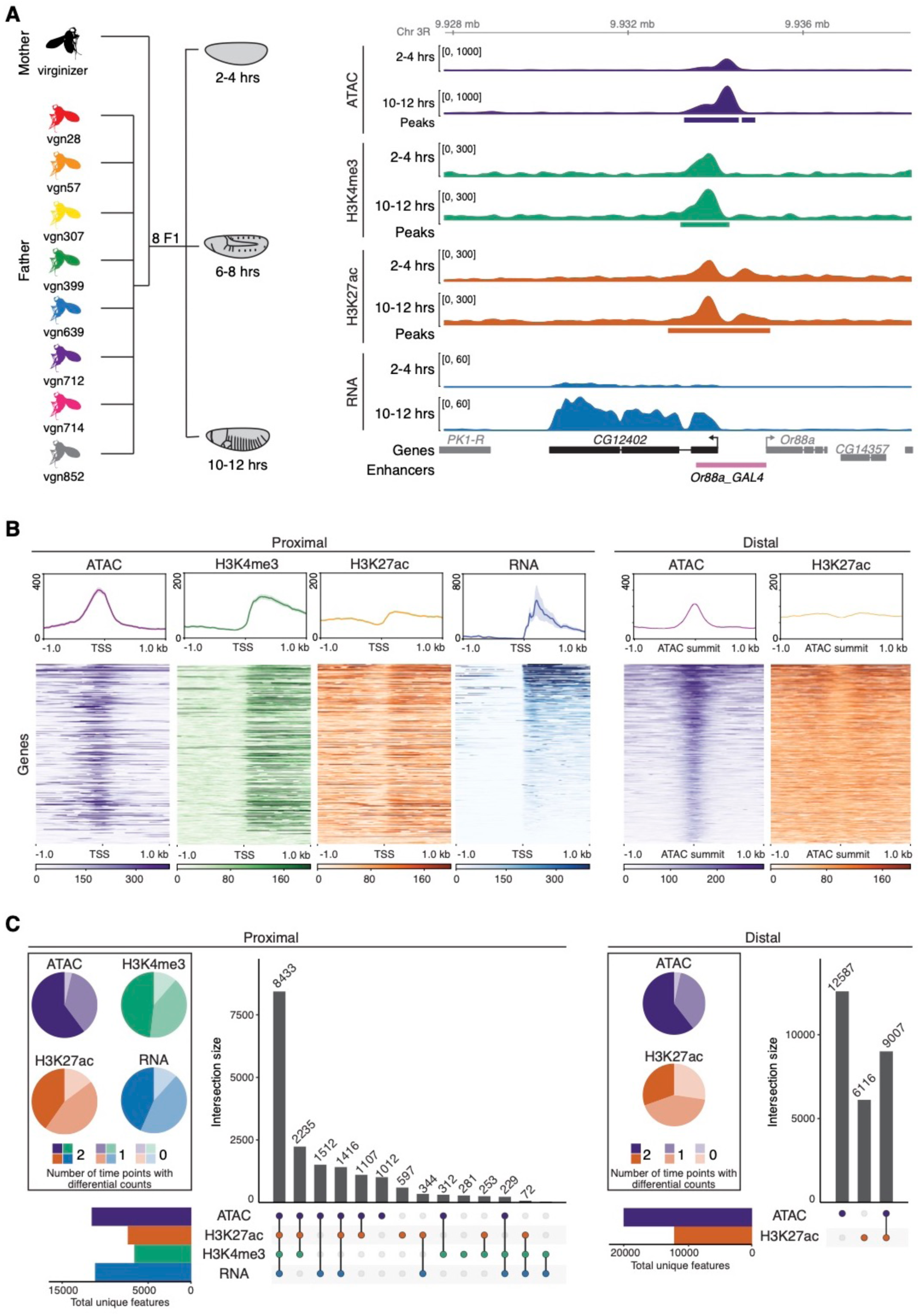
Quantifying gene expression and regulatory element activity in hybrid embryos. **a**. Left: Experimental design and data structure. RNA-seq, ATAC-seq and iChIP of H3K4me and H3K27ac were performed on embryos of three developmental stages from 8 F1 hybrids with a common maternal line. Right: Genome browser overview for the *CG12402* gene locus showing all data for 2-4 hours and 10-12 hours for the genotype vgn28. Bottom track shows characterized enhancers (Kvon et al. 2014). **b**. Top panel shows density plots for read count signal from each data type for TSS proximal and distal regions (left and right, respectively). Shaded regions indicate the 95% confidence intervals. Plots are centered at the TSS for promoter proximal regions, and ATAC summits for distal regions. Bottom panel shows a heatmap representation of the data type corresponding to the density plots shown above where rows are sorted by mean RNA-seq and mean ATAC-seq signal. **c**. Upset plots show the colocalization of signal for proximal and distal regions (at peaks in regulatory regions and genes) for all four data types. Regions common between data types (filled circle) are joined by a vertical bar. Horizontal bar plots indicate the number of unique genes/features. Pie charts show the proportion of features with statistically different total read counts between time points (color indicates the number of times (0/1/2) the feature is differentially expressed).

The F1 embryos were collected at three important stages of embryogenesis: 2-4 hours after egg laying, consisting primarily of pre-gastrulation, unspecified embryos (mainly stage 5), 6-8 hours (mainly stage 11), when major lineages within the three germ-layers are specified, and 10-12 hours (mainly stage 13), during terminal differentiation of tissue lineages (Fig. 1a). For each developmental stage, RNA-Seq, ATAC-Seq, and iChIP for H3K27ac and H3K4me3 (Buenrostro et al. 2013; Lara-Astiaso et al. 2014) were performed from the same collection of embryos (4 measurements x 3 stages x 8 genotypes = 96 samples). In addition, we collected samples from the parents of one F1 genotype, forming a parent/offspring trio that allowed us to partition genetic differences between the parents into *cis* and *trans* (Wittkopp et al. 2004). All measurements were made in replicates from independent embryo collections to assess biological and technical variability, giving a total of 240 samples (192 F1 samples (96 x 2 replicates) + 48 parental (4 measurement x 3 stages x 2 genotypes x 2 replicates)). Read counts were highly correlated between biological replicates, with median correlation coefficients of 0.98 for RNA, ATAC and histone data (Fig. S1b, Methods).

To define non-coding features, ATAC-Seq and ChIP-Seq reads from each cross were mapped to each parental line independently and the significant peaks merged to produce a combined set of common peaks used in subsequent comparisons across all genotypes. In total, we identified 11,211 genes with detectable expression, 31,963 ATAC-Seq peaks, 19,769 H3K27ac peaks, and 6,648 H3K4me3 peaks, active at one or more stages of embryogenesis (Table S2). Of these, 93.9%, 95.8%, 95.2%, and 96.9%, respectively, contained at least one SNP that distinguishes maternal and paternal haplotypes in at least one line. The *CG12402* locus, a predicted ubiquitin-protein transferase, provides a good example of overall signal quality (Fig 1a). The gene has dynamic expression, transitioning from very low to high expression from 2-4 hours to 10-12 hours (Fig. 1a, RNA-seq), which is accompanied by quantitative changes in chromatin accessibility, and to a lesser extent in histone modifications in its promoter-proximal region.

To examine the regulatory relationships between these different signals, we divided the data into promoter proximal (within +/- 500 bp of an annotated transcriptional start site (TSS) or H3K4me3 peak) or distal (putative enhancer) elements. Looking globally at promoter proximal regions, all signals show the expected enrichment and distribution around the TSS (Fig 1b, proximal), demonstrating the quality of the data. The ATAC-seq signal, for example, is highest directly at the promoter, representing occupancy of the basal transcriptional machinery, while H3K27ac and H3K4me4 signals are highest at the +1 nucleosome, reflecting the predominantly unidirectional nature of *Drosophila* promoters (Core et al. 2012; Mikhaylichenko et al. 2018). Moreover, H3K27ac has the expected higher signal levels at promoters compared to distal sites, (Kheradpour et al. 2013; Kwasnieski et al. 2014). Interestingly, while all three regulatory signals (ATAC-seq, H3K27ac and H3K4me3) are highly correlated at the promoters of actively transcribed genes (8,433 promoters contain all 4 signals, Fig 1c, left upset plot), 3,907 regions marked by H3K4me3 and overlapping peaks of ATAC-seq and/or H3K27ac show no detectable RNA-signal (Fig. 1c bar plots, 1b). Approximately 850 of these involve annotated transcripts of non-coding RNA (Flybase annotation) that lack a poly-A tail and were thus not selected in our Poly-A+ RNA-seq library. This suggests a surprising number of additional unannotated transcriptional events even within the well-annotated *Drosophila* genome.

The majority of H3K27ac (62.5%) and ATAC peaks (63.7%) are distal to an annotated promoter, and likely represent enhancer elements. Of the distal ATAC peaks, 58% (12,587/21,594) have no H3K27ac signal and may represent inactive enhancers or other regulatory elements, e.g. insulators (Fig. 1c). The remaining 9,007 distal elements overlap H3K27ac signal (Fig. 1c, right), which is generally bimodally distributed around the ATAC-seq peak (Fig. 1b), suggestive of active enhancers. Conversely, 40% (6,116/15,123) of H3K27ac peaks do not overlap a significant ATAC peak (Fig. 1c, right) and represent regions with quantitively lower ATAC signal (Fig. S1c), below our stringent threshold for detection. Both gene expression (RNA-seq) and non-coding elements (based on ATAC-seq and chromatin signatures) show evidence of dynamic activity, with the majority (72%-96%) of features showing statistically significant changes in total counts between developmental time points across all *F1* lines (Fig. 1c, pie charts; Methods), *CG12402* being one example (Fig. 1a).

Taken together, these features demonstrate both the quality and richness of the data and its usefulness to further annotate the regulatory landscape of the *Drosophila* genome at these important stages of embryogenesis.

### Allele-specific variation is common across genotypes and regulatory layers

To examine the impact of genetic variation, reads from each cross were mapped to personalized genomes for each parent and assigned to the maternal or paternal haplotype, where possible (Methods). To test for allele-specific differences for each gene per line and time combination, we used an empirical Bayes framework to model allele-specific counts for each data type using a beta-binomial model (Fig. S2a). Most promoter proximal and distal elements had the expected allelic ratios centered at 50:50 across autosomes (Fig 2a), with a slight elevation in the magnitude of allelic imbalance (AI) at distal sites (Fig. S2b). RNA allelic ratios were also concordant with the direction of change of embryonic eQTL (Fig. S2c), previously quantified in the same paternal lines at the same stages of embryogenesis (Cannavo et al. 2017), further verifying our approach.

**Figure 2:**
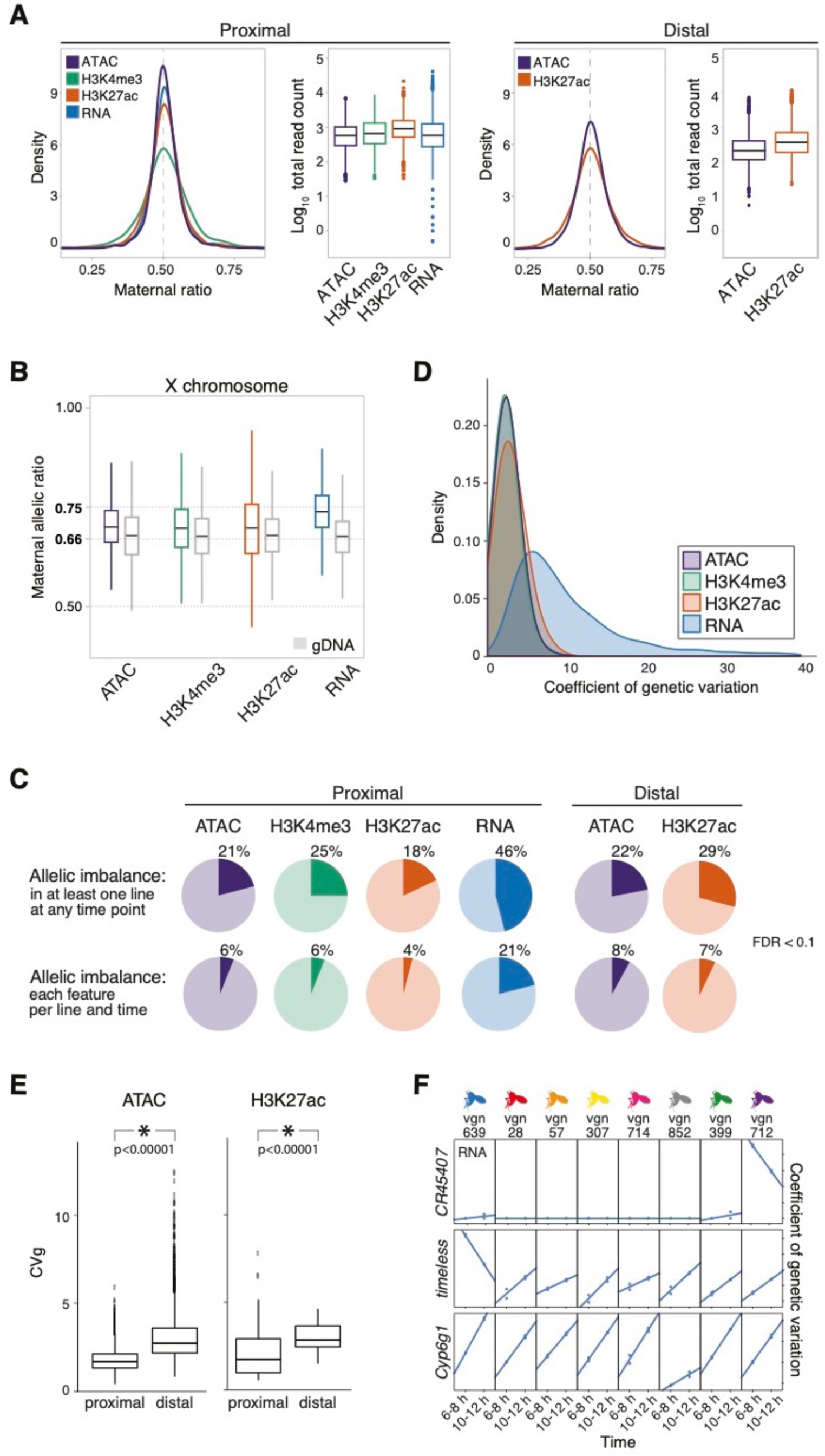
Allelic imbalance is common across regulatory data types. **a**. Density plot of allelic count distribution and matching boxplot showing total read count abundance (log_10_) in the autosomes at TSS proximal (left) and distal (right) regions for all data types assayed. **b**. Box plot shows the distribution of the maternal allelic ratio of X chromosome in each data type. Each distribution is compared to the allelic ratio observed in genomic DNA in grey. **c**. Pie charts show significantly allelic imbalance (AI) genes/features at promoter proximal (left, TSS +/-500 bp) and distal (right, 500-1500 bp +/- from TSS) regions for all four data types (FDR<0.1). *Upper*: AI events in at least one F1 line at any time point. *Lower*: AI events detected in all 8 F1 lines in all time points, on a per line and time basis. **d**. Smoothed histograms show the distribution of coefficients of genetic variation for all features with statistically significant between-line variances within each regulatory layer. **e**. Box plots show the distribution of coefficient of genetic variation (CVg, y axis) for chromatin accessibility (left) and H3K27ac signal (right), for promoter -proximal and -distal sites. Genetic influences are more pronounced at distal regulatory elements in ATAC and H3K27ac. **f**. Line plots show three examples of individual lines having distinct expression profiles. Coefficients of genetic variation are typically larger for RNA than for non-coding features, an effect that often results from one or two lines having significantly altered expression relative to the panel as a whole.

To evaluate sex ratios in the embryo pools, and to set a reference point for evaluating allelic imbalance and dosage compensation on the X-chromosome (Lucchesi and Kuroda 2015), we sequenced the genomic DNA (gDNA) of each cross. This confirmed that our embryonic pools were relatively sex balanced, with the expected X-chromosome allelic ratio of ∼0.66 observed across our gDNA dataset (Fig 2b). Consistent with full dosage compensation on the maternally-derived male X chromosome (Georgiev et al. 2011), we observed a maternal:paternal ratio of 0.74 for RNA (Fig. 2b; Methods). Interestingly, a similar degree of up-regulation (dosage compensation) was not observed for chromatin data: for both chromatin accessibility and histone modifications, the observed ratio at X chromosome sites (H3K27ac=0.688, H3K4me3=0.692) is more similar to the observed genomic ratio of 0.66 than to the expected ratio of 0.75 under full dosage compensation (Fig. 2b). The ratios showed no significant difference when comparing proximal to distal sites, arguing against the hypothesis that the two-fold upregulation of gene expression on the male *Drosophila* X chromosome results from a two-fold increase in the loading of polymerase at its genes’ promoters (Conrad et al. 2012). Our results rather indicate that whatever the mechanism of dosage compensation in *Drosophila*, it does not lead to a linear increase in chromatin accessibility on the male X chromosome, though some increase in accessibility on the upregulated X is consistent with our measurements (Urban et al. 2017; Pal et al. 2019). Regardless of its cause, we used the empirically observed average ratio for X-chromosome features for each data type to form the null-hypothesis in subsequent beta-binomial tests for allelic imbalance.

Overall, allelic imbalance is common, with 46% of genes and between 18-25% of non-coding features showing statistically significant AI in at least one line at one or more time point (Fig. 2c, FDR <0.1). The magnitude of AI is generally evenly distributed across SNPs with a range of minor allelic frequencies, however highly imbalanced peaks show a strong enrichment for extremely rare SNPs (including potentially *de-novo* mutations) found uniquely in the maternal line relative to the 205 lines of the full DGRP panel (Fig. S2d, Chi^2 test; p<2.2e-16), highlighting the disproportionate impact of rare and *de-novo* mutations on expression phenotypes (Cannavo et al. 2017). Allelic imbalance is more frequently observed for RNA than for other regulatory layers (Fig. 2c). In contrast to what is observed in mammals (Villar et al. 2015), promoter-proximal elements are slightly more polymorphic (pair-wise differences (pi) = 0.132 vs 0.129, Wilcoxon-test p = 1e-10) and evolve faster (phyloP = 0.514 vs 0.560, Wilcoxon-test, p < 2.2e-16) in *Drosophila* as compared to distal elements (putative enhancers) (Table S3). Despite this, distal peaks of open chromatin and H3K27ac show larger (Tukey’s ASD, p < 0.0001) and more frequent (χ^2^ test, p < 2.2e-16) allelic imbalance than their proximal counterparts (Fig S2b).

To understand how allelic-imbalance relates to heritable variation at the total count level, we took advantage of the fact that our measured F1 lines share a common maternal genotype. As a result, line effects (from a linear model) are expected to be directly proportional to heritability – the degree to which phenotypic variation can be explained by genetic factors (Lynch and Walsh 1998). To make these effects comparable across genes and features, line effects (standard deviations) were scaled by the mean read count of each feature, expressing line effects as a percentage deviation from the mean phenotype (coefficient of genetic variation). For chromatin features, the magnitude of genetic variation on measured signal is relatively modest, with the average peak varying by ∼5-10% of the mean phenotype among crosses (Fig. 2d). Overall, the effects of heritability genetic variation is higher at distal regulatory elements as compared to their proximal counterparts (Fig. 2e, p < 1e-5, Methods), consistent with the greater magnitude of AI at distal sites. For RNA, in contrast, the magnitude of genetic effects is pronounced, with an average coefficient of genetic variation of ∼9% and some genes showing coefficients of ∼40%, indicating that genetically encoded differences in expression can account for nearly half of some genes’ mean expression levels. In many cases, such high coefficients are driven by one or a few lines showing highly divergent patterns of expression (Fig. 2f, right) suggesting that large-effect gene expression differences in this population can be driven by large-effect *cis*-acting mutations.

### Allelic imbalance is pronounced in metabolism and environmental response genes

Imbalanced genes and associated regulatory features are enriched for fast-evolving and *Drosophila*-specific genes lacking clear categorical annotations (Mi et al. 2003; Turner et al. 2008) and are depleted in TFs and their associated regulatory elements (Fig. S3, Table S4), consistent with our previous eQTL study (Cannavo et al. 2017). AI is also enriched for metabolic genes at the RNA level, although interestingly this is not observed for associated regions of open chromatin or histone modifications (Fig. S3, Table S4). The observed differences in AI among gene categories may reflect differential histories of selection; regulatory regions in the vicinity of TFs show a depletion of nucleotide diversity (pi, rank biserial correlation = −0.052, p < 1e-4) and harbor more low-frequency SNPs (rank biserial correlation = −0.173, p = 2.8e-3; Table S4) compared to background. However, this difference AI could also be explained by different gene categories having different sensitivities to mutations (buffering), a point we explore further below.

For most gene categories, AI is equally likely to favor the maternal or the paternal allele. However, a subset of categories shows consistent and often large parent-specific biases, a trend that is particularly striking for male-biased genes associated with immunity or insecticide resistance (Fig. S4a; Table S5). *Cyp6g1*, for example, is not expressed in embryos of our maternal line (Fig. S4b), which is derived from a laboratory stock isolated before the widespread use of agricultural pesticides, or in embryos sequenced by the modENCODE project (Celniker et al. 2009). It is, however, strongly upregulated in every measured paternal haplotype from the wild, and its overexpression contributes to DDT resistance in multiple *Drosophila* species (Fig. S4b, (Daborn et al. 2001; Battlay et al. 2016)). Highly imbalanced genes like *Cyp6g1* (Table S5) overlap genes whose expression varies extensively among lines (Fig. S4c, p < 1e-6) and who have high levels of heritability, highlighting the important contribution of selection on *cis*-regulatory elements in shaping responses to changing environments.

### The impact of cis-acting genetic variation is largely consistent across development

We next evaluated if, and to what extent, allelic ratios change during development. Overall, we observed a surprising constancy of allelic imbalance between embryonic time points: Despite the temporal modularity of many *cis*-regulatory elements, imbalanced features at one time point have a ∼50% chance of being imbalanced in the subsequent time-point (Fig. 3a, S5a). To further quantify the potential impact of development on allelic ratios, we constructed a series of linear models comparing the effect sizes of genetics (genotype/line effect) vs developmental stages (time effect) on total counts and allelic ratios across our experimental design. For total counts, developmental time was the greatest contributor to variation across all data types (Fig. 3b, upper panel), consistent with the clear time specific clustering by principal component analysis (Fig. 3b, lower panel, shown for RNA, Methods). Interestingly, this predominance of time is largely restricted to distal and not proximal sites for ATAC-Seq (Fig 3d), likely reflecting the frequently constitutive accessibility of promoters during *Drosophila* embryogenesis (Cusanovich et al. 2018). In contrast to the total counts, the impact of developmental time on allelic ratios is significantly reduced compared to genetic (line) effects (Fig 3c, upper panel). Correspondingly, there is a lack of time-point specific clustering in PCA (Fig 3c, lower panel), although there are some examples of allelic ratios that change over time in a coordinated manner between regulatory layers (Fig. S5b).

**Figure 3:**
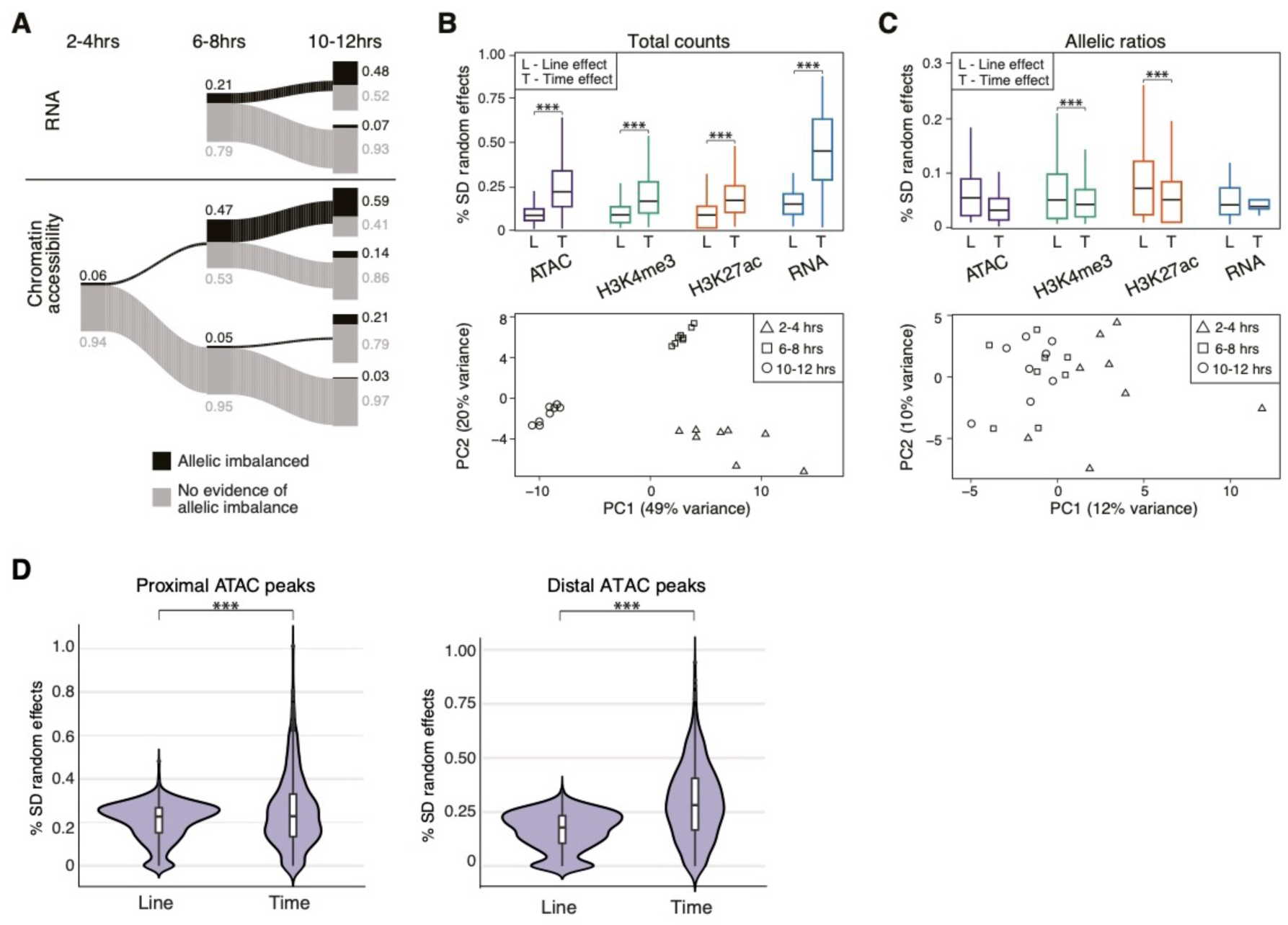
Allelic imbalance is generally not predictive of developmental time. **a**. The relationship of allelic imbalance across time points for RNA (upper panel) and chromatin accessibility (lower panel). Proportions of AI and non-AI features are shown in black and grey, respectively, and represented by the thickness of line. Exact proportions for each category are provided as numbers. Data for 2-4 hour time point for RNA are not included due to presence of maternal transcript at this stage. **b**. *Top*: Box plots show the distribution of effect sizes obtained from mixed linear models, for total counts. For each type of data (gene/feature), the effect sizes of time (T) and line (L) effects are shown. *Bottom*: Principal component analysis of gene expression for total counts. **c**. *Top*: Box plots showing the distribution of effect sizes obtained from mixed linear models, for allelic ratios. For each type of data (gene/feature), the effect sizes of time (T) and line (L) effects are shown. *Bottom*: Principal component analysis of gene expression for allelic ratios. **d**. Results from mixed linear models examining the effect of developmental time versus line (genotype) between proximal and distal ATAC-seq peaks for total count data. Distal peaks show a larger time effect compared to genotype effect (Mann-Whitney U, p<2.2e-61). This is only slightly evident for promoter proximal peaks (Mann-Whitney U, p<8.5e-5).

Interactions between genetic and developmental effects can play an important role in gene regulation (Paaby and Gibson 2016; Yadav et al. 2016). We therefore looked for evidence of interaction effects in linear models fitted to total counts or allelic ratios containing only time, only genotype, time plus genotype (time + genotype), or interactions between the two (genotype x time (GxT)). Interaction effects occur frequently at the total count level and are particularly common for gene expression, making up nearly 30% of all analyzed models (Table S6) and highlighting a potentially important role for developmental stage by genetic (GxT) interactions in population-level variation during embryogenesis, as previously observed (Cannavo et al. 2017). In contrast, there is little evidence for interaction effects at the level of allelic ratios for gene expression or ATAC-seq peaks (Table S6), consistent with the relatively small numbers of allelic ratios reported to show influences of gene x environment interactions across environmental conditions (Moyerbrailean et al. 2016; Knowles et al. 2017).

In summary, allelic effects are often larger at distal compared to promoter regions, with allelic effects at both regions being surprisingly stable across develompental time points. In contrast, total counts vary dramatically between time points, with interactions between genotype and developmental stage (GxT) being common, particularly for gene expression. Given that total counts are influenced by genetic variation in both *cis* and *trans*, this highlights an important role for *trans* acting variation in the maintenance of evolutionarily relevant interaction effects

### Information flows in different directions across cis-regulatory layers

Although quantitative signals at chromatin features are highly correlated with gene expression, the relative causal relationships between chromatin accessibility, histone modifications, and gene expression remain unclear. To assess this, we used allelic ratios as a perturbation measured at different regulatory layers to model the paths by which genetic variation influences regulatory phenotypes. Allelic ratios in all pairs of datatypes are correlated, to varying extents (Fig. 4a), and in all cases we could reject the null hypothesis of independence (e.g. highest p-value between all comparisons was 4.2e-17 for ATAC and H3K4me3). We tested for an enrichment/depletion of co-occurring, statistically significantly imbalanced (FDR < 0.1) genes/features using an intersection-union test (Fig 4b; Methods), using a distance of +/-1500 kb to assig distal features to genes. The co-occurrence of allelic imbalance is especially pronounced for chromatin features, in particular H3K4me3 and proximal H3K27ac with a log-odds >2.0 (Fig. 4b). Interestingly, for chromatin accessibility and H3K27ac, the co-occurrence of AI is more pronounced at promoters (proximal) than enhancers (distal) (Fig. 4b), despite allelic imbalance being slightly more frequent (p<2.2e-16, Fig. 2c) and of greater magnitude (Fig. S2b) at distal sites. This suggests that H3K27ac and chromatin accessibility are more functionally coupled at promoters compared to enhancers, perhaps reflecting the fact that not all active enhancers seem to require H3K27ac (Bonn et al. 2012; Pradeepa et al. 2016).

**Figure 4:**
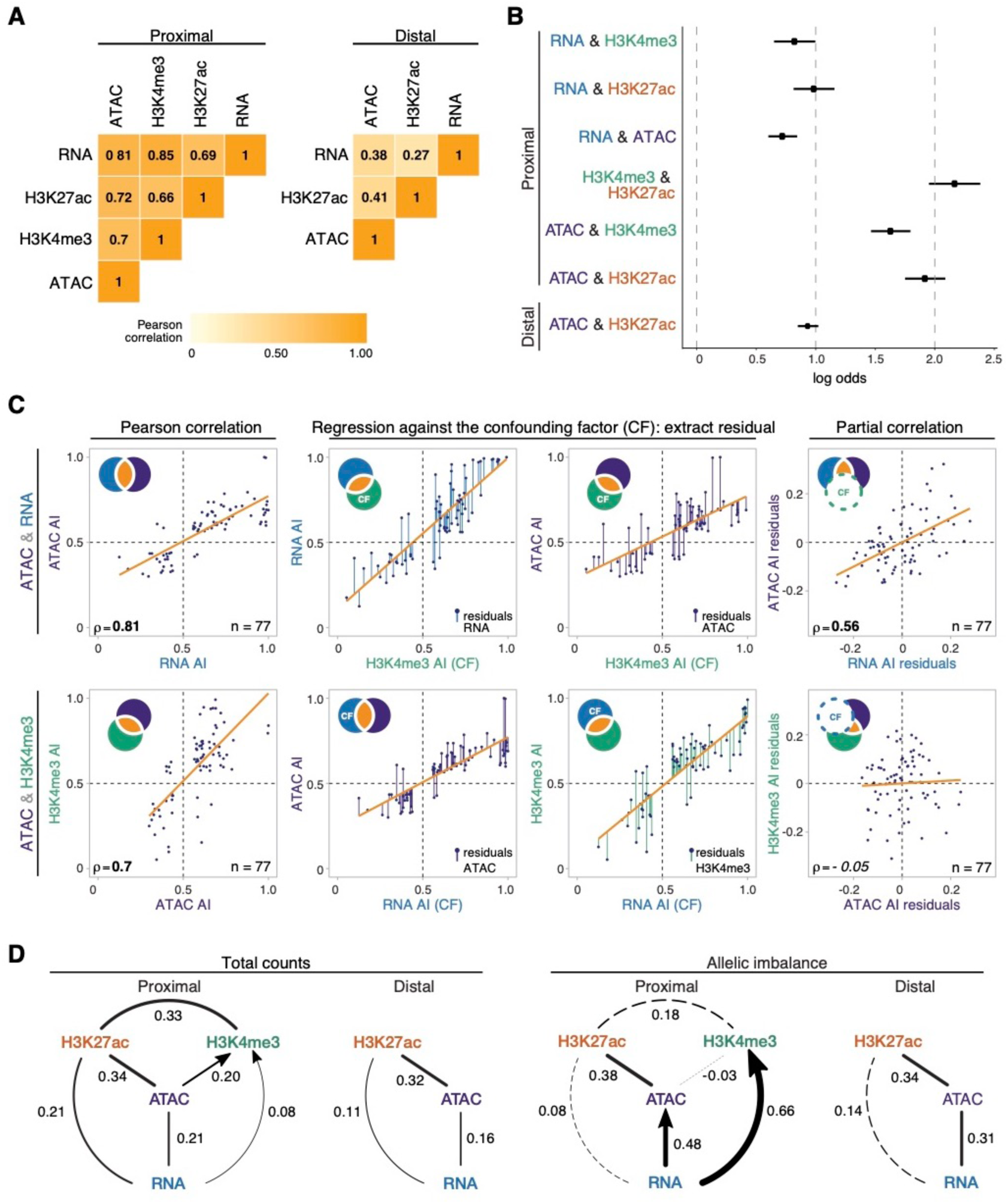
Allelic imbalance is propagated through regulatory layers via different epigenetic paths. **a**. Heatmaps show Pearson correlation coefficient of allelic ratios between each pair of data type for promoter proximal (left) and promoter distal (right) regions. Data restricted to 6-8 hours plus 10-12 hours, and features/genes whose allelic ratio exceeds 0.5 +/- 0.06. **b**. Increased log odds of co-occurrence of allelic imbalance between two regulatory layers. X-axis shows the log odds based on intersection-union tests (Methods). 6-8 hours plus 10-12 hours data is shown. Bars stemming from dots are 95% confidence intervals. **c**. Stepwise example of a partial correlation analysis of allelic ratios for three variables (ATAC, RNA and H3K4me3). Partial correlation analysis is shown between gene expression and chromatin accessibility (upper row), and promoter proximal H3K4me3 and chromatin accessibility (lower row). Venn diagram schematics (top left) illustrates the variance of each variable and its shared proportion (orange), as measured by linear regression (orange lines). *Left panels*: Pearson correlations for the two comparisons are significant. *Middle panels*: regression of each initial variable against a third, confounding variable (H3K4me3, upper row; RNA, lower row). Residuals of the initial variables (colored lines) represent the non-overlapping part of the circle of the same color in the schematic. *Right panels*: correlations of the residuals, which exclude the variance shared by the confounding factor (dashed circle in schematic). This resulting partial correlation is not significant in the bottom example, suggesting a lack of direct correlation within the pair H3K4me3-ATAC. **d**. Partial correlation and directional dependency regression for total counts (left) and allelic ratios (right). Significant partial correlations (solid lines) suggest dependencies among regulatory layers. For each significant edge (p < 0.01), copula regression was used to assign directionality (arrows, delta > 0.01). Line thickness indicates the value of partial correlations, dashed lines indicate non-significance. Results for promoter proximal and distal regions shown separately.

Due to the large amount of covariation between the different regulatory features (Fig. 4a), it is difficult to infer causal relationships from correlation data alone. To address this, we used partial correlation to identify independent, pairwise correlations between multiple co-varying variables beyond their global correlations after thresholding on allelic ratios to remove features/genes with low information content (Fig. 4c, S6a, Methods) (Lasserre et al. 2013; Pai et al. 2015). We first analyzed the total count data to evaluate the overall relationships among histone modifications and gene expression. Our results closely mirror those of Lasserre *et al* in CD4+ and IMR90 cells (Lasserre et al. 2013), including finding a clear relationship between gene expression levels and the total abundance of H3K27ac that ‘explains away’ (at least in a statistical sense) much of the correlation between gene expression and promoter-proximal H3K4me3 (Fig. 4d, left). We also observed a statistically significant relationship between open chromatin and gene expression, though the strength of this partial correlation is reduced relative to standard Pearson correlation analyses (Fig. S6b).

To assess the functional impact of *cis*-regulatory perturbations, we next applied the partial correlations analysis to allelic ratios (Fig. 4d, right). Relative to the total count data, allelic ratios reveal a much stronger relationship between open chromatin and gene expression for both proximal and distal regulatory elements (Fig. 4d, right), highlighting an important, possibly causal, link between mutations affecting accessibility (presumably TF occupancy) and gene expression. A significant correlation is also observed between H3K27ac and open chromatin at promoters, though interestingly, we see little evidence for a direct relationship between H3K27ac and gene expression itself (Fig. 4d, right). The latter is surprising as it differs from what is observed with total count data, and suggests that although promoter H3K27ac is highly correlated with, and even predictive of gene expression (Karlic et al. 2010), they may not be mechanistically directly linked. In contrast, allelic ratios for promoter proximal H3K4me3 show strong evidence of a direct correlation with gene expression that is independent of, at least in a statistical sense, allelic differences in chromatin accessibility or H3K27ac (Fig. 4d, right). Taken together, this analysis suggests two independent pathways by which selection on segregating mutations could influence gene expression, one affecting open chromatin and promoter-proximal H3K27ac, and the other influencing H3K4me3.

To explore these relationships further, we analyzed each edge identified by partial correlations using copula directional dependence analysis (Kim et al. 2008; Lee and Kim 2019), a statistical approach based on copula regression that evaluates the directionality of the pairwise relationships allowing for non-linearities (Methods). This method assigns a direction to each edge for which there is clear evidence for greater explanatory weight in one direction. For TSS-proximal regions, this placed RNA upstream of both H3K4me3 and open chromatin (Fig. 4d right, arrow). Although counter intuitive at first glance, this suggests that gene expression is not highly sensitive to variations in H3K4me3, while conversely changes to RNA is more predictive of H3K4me3 enrichment. This could reflect redundancy between regulatory elements, i.e. changes in a single open chromatin region, as tested here, may not be sufficient to impact expression in an allele-specific manner. This result is also consistent with the hypothesis that H3K4me3 is not functionally required for transcription, but may rather be deposited as a consequence, and be involved in post-transcriptional events, as recently proposed (Howe et al. 2017). Similarly, allele-specific variation in RNA better explains variation in chromatin accessibility compared to the reverse, i.e. not all variation in open chromatin leads to a corresponding change in gene expression (Fig. 4d, right).

In summary, *cis*-acting genetic variation shows greater covariance between open chromatin and H3K27ac enrichment at promoters compared to putative enhancers. By measuring informative dependencies on the impact of *cis*-acting genetic variation, we identified multiple epigenetic pathways affecting transcription. Specifically, genetic variation acts to change gene expression levels via the interplay between at least two different promoter-proximal paths: open chromatin and H3K27ac, or H3K4me3. Moreover, the flow of information suggests that gene expression is often buffered against *cis*-acting mutations (presumably affecting TF binding) at associated regulatory elements.

### Regulatory buffering varies depending on gene function and local chromatin architecture

Genes from different functional categories often have differences in the complexity of their regulatory landscapes. Metabolic genes, for example, typically have relatively simple and more compact regulatory landscapes with fewer enhancers that are generally located close to the gene’s promoter (Zabidi et al. 2015). TFs, in contrast, have many enhancers often with partially overlapping spatial activity (“shadow enhancers”) that are located at varying distances from the gene’s promoter (Spitz and Furlong 2012; Long et al. 2016). This additional regulatory complexity is thought to make TFs more robust to deletions and sequence variation affecting their regulatory elements (Xiong et al. 2002; Cretekos et al. 2008; Montavon et al. 2011; Cannavo et al. 2016). To examine this, we assessed the extent to which allelic imbalance in the expression of different gene categories is independent of, or decoupled from, imbalance in their associated regulatory elements, treating gene categories with greater independence as ‘buffered’. Among all comparisons of conditional probabilities, the expression of TFs, transmembrane genes, ancient genes (conserved bilaterian processes), signalling pathway genes, secreted genes, are most insensitive to imbalance in other regulatory layers (Fig. 5a). In contrast, genes and their regulatory elements associated with cytoskeletal function, glycoproteins, and notably metabolism show an increased sensitivity to allelic imbalance in other regulatory layers (Fig. 5a). Taken together, our analyses suggest that in addition to purifying selection acting to remove genetic variation, regulatory buffering helps to ensure robust expression of TFs and other developmental regulatory factors from the effects of *cis-*acting mutations.

**Figure 5:**
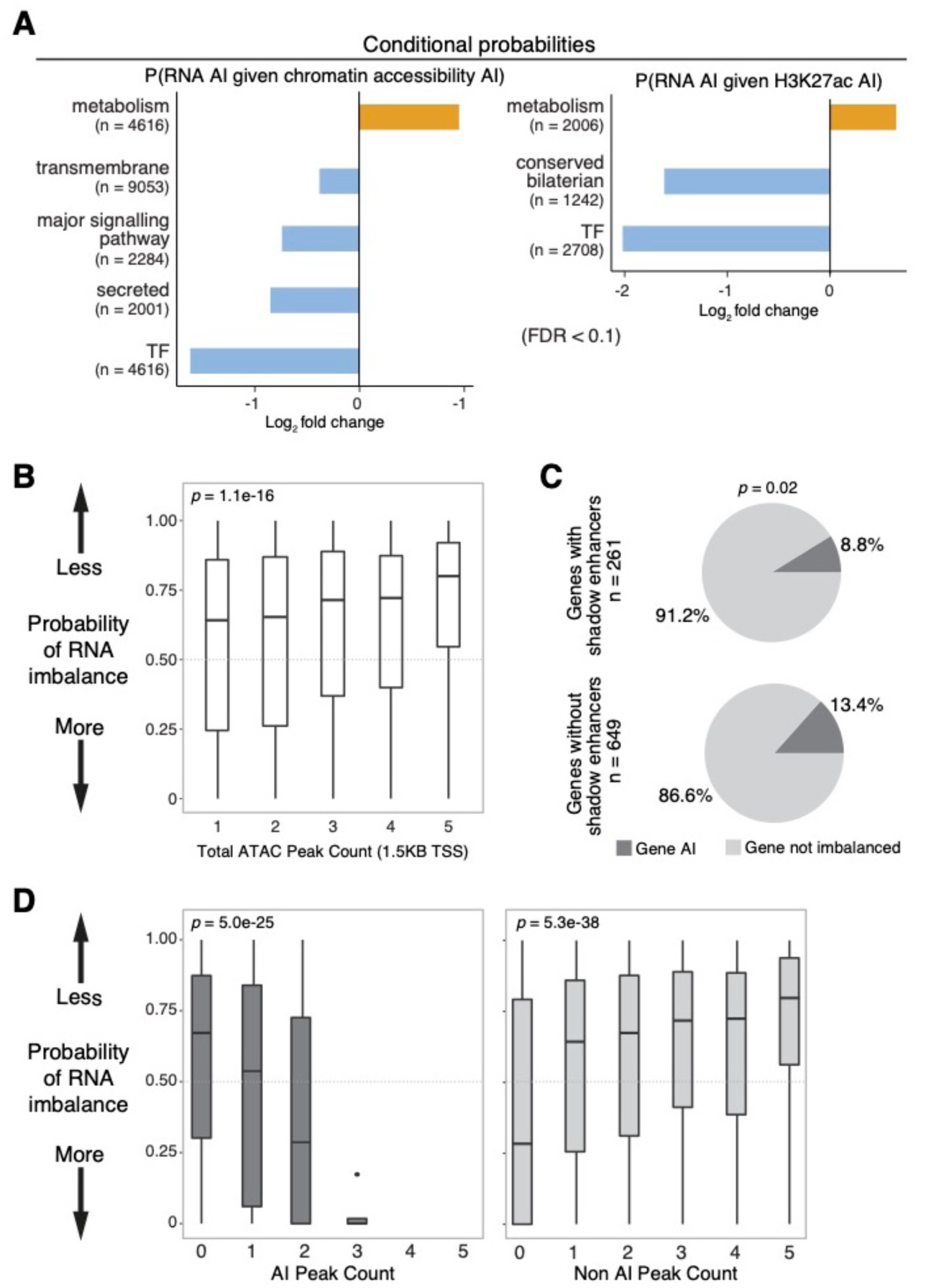
Regulatory buffering varies across gene categories and with local chromatin structure. **a**. Conditional probability of allelic imbalance in gene expression given allelic imbalance in associated chromatin peaks (left) and regions of H3K27ac (right) across gene categories. X-axis show log2 fold change where background is based on genome-wide expectation. Gene categories enriched (orange) or depleted (blue) for imbalance, relative to background, are indicated (FDR>0.1, Fisher’s exact test). **b**. Box plots denote the probability of allelic imbalance in gene expression based on numbers of neighbouring ATAC peaks (TSS < 1.5kb). Genes associated to more ATAC peaks are more likely to show similar expression in both alleles compared to genes with fewer peaks. **c**. Pie charts displaying the proportion of genes with allelic imbalance in RNA associated to ATAC-seq peaks overlapping known partially redundant/shadow enhancers (top) or not (bottom). Genes associated with shadow enhancers are less likely to be allelic imbalanced compared to genes without (*x*^2^=5.3, p=0.02). **d**. ATAC-seq peaks have a cumulative effect on gene expression. The probability of imbalance in gene expression (y-axis) is shown as a function of the number of ATAC-seq peaks that are allelic imbalanced (left) or not imbalanced (right).

To more directly assess the relationship between buffering and regulatory complexity, we compared the number of ATAC-seq peaks in a gene’s regulatory domain (+/- 1.5kb TSS) with the probability of imbalance in that gene’s expression. Imbalanced genes have fewer associated ATAC-seq peaks genome-wide (Kruskal Wallis p =1.1e-16, Fig 5b). This trend is particularly striking for single-peak genes, which have significantly more AI than genes with multiple associated regulatory elements (Mann-Whitney U test, p-value=6.4e-6). Consistent with the observation of transcriptional robustness (a lack of AI) for genes with multiple regulatory elements, genes associated with partially redundant enhancers (or shadow enhancers) have a modest reduction in the frequency of allelic imbalance compared to genes without (Fig. 5c, *x*^2^=5.3, p=0.02), based on a previously defined set of shadow enhancers for mesodermal genes (Cannavo et al. 2016). We note, however, that this buffering is not absolute – even genes with multiple regulatory elements are more likely to be imbalanced when multiple associated peaks show unbalanced allelic ratios (Fig. 5d).

In summary, there is a relationship between a gene’s regulatory complexity and the degree to which its expression is influenced by non-coding genetic variation in its regulatory elements, with more regulatory elements providing a degree of buffering against genetic perturbations. Allelic imbalance at multiple enhancers in the vicinity of a gene can have a cumulative influence on allele-specific gene expression.

### Variation in gene expression is less heritable than for chromatin features

Gene expression phenotypes are influenced by linked *cis*-acting genetic variation, but also by *trans* acting variation that is not directly captured using F1s alone. To estimate the relative impact of *trans*-acting variation, we performed the same experiments (iChIP-Seq, ATAC-Seq, and RNA-Seq) on a trio of lines consisting of one F1 line and stage-matched embryos from the maternal (vgn) and paternal (DGRP-399) lines. As the two homologous chromosomes in F1 cells have a common nuclear *trans* environment, allelic ratios in F1s estimate *cis*-based differences between the two parents. Differences in parental read counts not reflected in F1 allelic-ratios give an estimate of *trans*-acting contributions to between line divergence (Landry et al. 2005; Tirosh et al. 2009; Goncalves et al. 2012; Wong et al. 2017).

Using a maximum likelihood framework, we classified features as *cis, trans, cistrans*, or *conserved* and found a similar distribution among categories for all non-coding chromatin features, with *cis*-acting effects being more common than *trans* (59% vs. 41%, p < 2.2e-16, chi^2; Fig. 6a, Methods, Table S7). This enrichment is particularly pronounced for histone modifications, with nearly twice as many *cis* influenced peaks compared to *trans* (Fig 6a, S7a). Gene expression, in contrast, is more strongly influenced by *trans*-acting genetic variation (Fig. 6a: 55% trans vs. 45% cis, p = 0.0073, chi^2). Moreover, a higher fraction of *cistrans* genes have more *trans* compared to *cis* variation, a pattern not observed for chromatin features (*trans* proportions 0.67 vs. 0.53, p = 2.77e-05; Fig. S7a).

**Figure 6:**
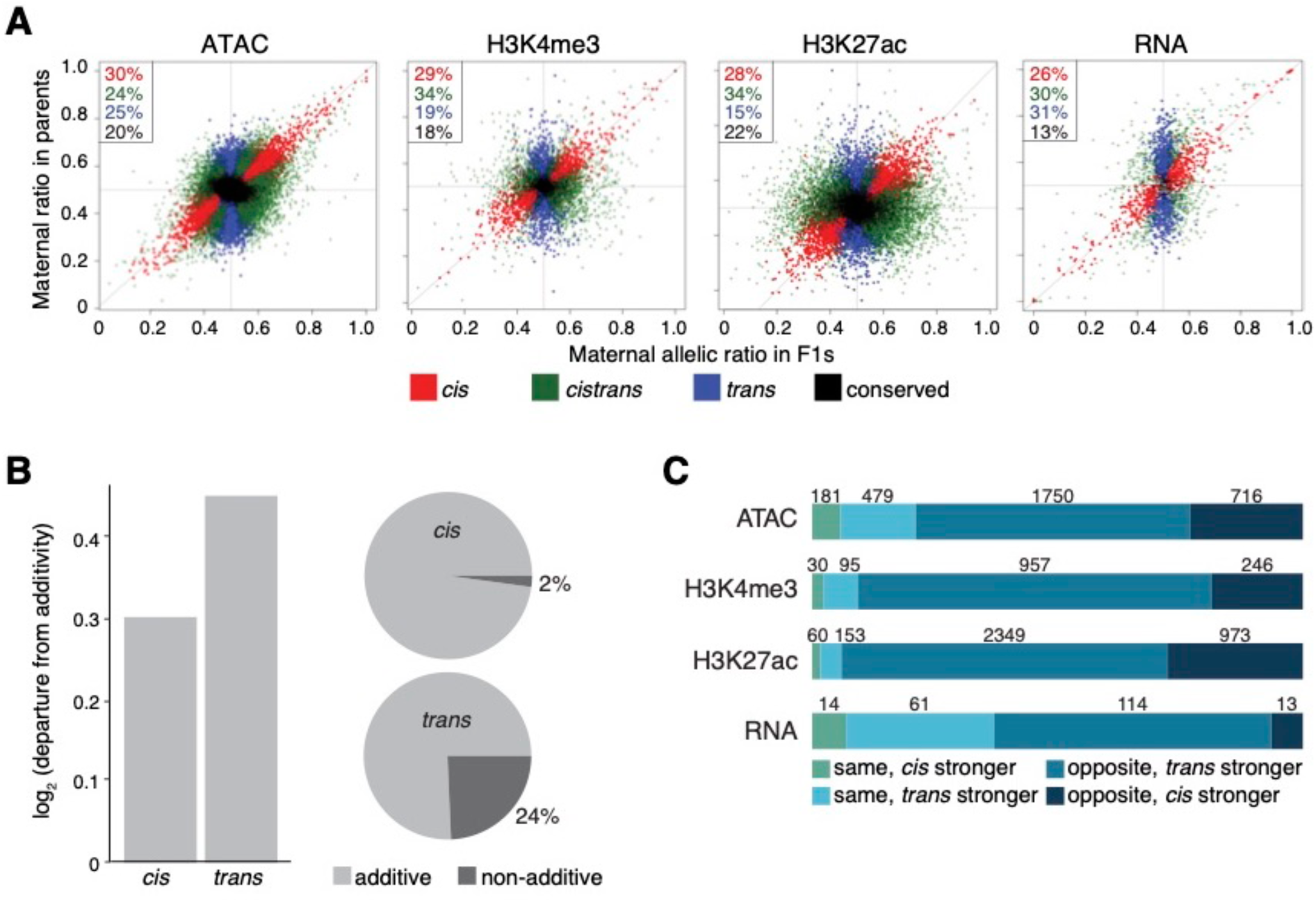
Chromatin features are more heritable than gene expression. **a**. Scatter plots of F1 allelic ratio (x-axis) against the maternal/paternal ratio observed in (normalized) parental, total count libraries. Genes/features along the diagonal are exclusively influenced by *cis*-acting variation, while vertically distributed genes/features show exclusively *trans*-influences. Colors indicate maximum likelihood classification into *cis, trans*, and *cistrans* (a mixture of *cis* and *trans*) or *conserved* genes/features. **b**. Left, bar plots shows the magnitude of deviation from additivity (parental mean) for features classified as *cis* vs. *tran* (BIC >= 2). Right, pie charts showing fraction of additive and non-additive genes for *cis* (upper) and *trans* (lower) classes. **c**. Classification of *cistrans* effects (BIC >= 2) for each regulatory layer into categories reflective of likely selective effects. Numbers and horizontal bars represent the size and relative proportions of each *cistrans* relative direction class in each data type. There is more directional selection (same directions, *cis* + *trans* > *cis*) than compensatory evolution (opposite directions, *cis* + *trans* < *cis*) in gene expression as compared to in chromatin features.

Previous studies suggest that the effects of *cis*-acting mutations are more likely to be inherited in an additive manner, compared to *trans* influences (Lemos et al. 2008; Meiklejohn et al. 2014; Wong et al. 2017). This has important consequences, as it is typically *additive* genetic variation that is most directly acted upon by natural selection (Lynch and Walsh 1998). We evaluated this in our data by examining the extent to which the F1 signal (total read count) for each gene/feature departed from the parental average (a strictly additive model). For open chromatin, whether influenced by *cis* or *trans*, we could reject a non-additive model in fewer than 1% of cases (Fig. S7b), consistent with the finding that most variation affecting TF binding is inherited additively (Wong et al. 2017). For gene expression, in contrast, the additive model could be rejected for 32% of genes, with *trans* influenced genes departing from an additive model far more often than *cis* (Fig. 6b: 24% vs. 2%, p < 1e-4).

To better understand the factors that contribute to the proportion of *trans*-acting variation (and by association, non-additive heritability), we examined the contribution of regulatory complexity. Mirroring our buffering results, genes with more regulatory elements in their vicinity are more likely to be classified as *trans-*acting (*trans* = 2.58 peaks per gene vs. 1.9, p = 0.00094) and more likely to show non-additive inheritance (non-additively inherited genes = 2.19 peaks per gene vs. 1.82 peaks per gene, p = 1.4e-3). Similarly, there is a significant, though modest, enrichment of *trans* influences among TFs and a depletion among metabolic genes, two categories that are strongly distinguished in the complexity of their associated regulatory landscapes (Table S8). Correspondingly, among 80 tested gene categories, DNA-binding TFs (p = 3e-3) and interestingly mitochondrial associated genes (p = 2e-6) stand out as the two gene categories with statistically elevated frequencies of non-additive inheritance (Fisher’s exact test; Methods). Thus, while TFs generally show reduced genetic variation among lines and reduced allelic imbalance in gene expression (Fig. S3), they are still affected by *trans*-acting variants whose non-additive inheritance reduces the efficacy by which selection can alter gene expression differences among different lines.

Genes influenced by both *cis* and *trans* acting variants (*cistrans*) provide an opportunity to understand patterns of recent selection: In the face of compensatory evolution, *cis* and *trans* acting influences are more likely to work in opposite directions, while directional selection will be more likely to reinforce *cis* and *trans* effects acting in the same direction. Using the classification of *cis* and *trans* by Goncalves et al (Goncalves et al. 2012), we observed that *cis* and *trans* effects were much more likely to act in a compensatory manner as compared to gene expression: for chromatin accessibility and histone modifications, 13% of *cistrans* features were classified as *same* vs. 37% for RNA (Fig. 6c: p < 2.2e-16 chi^2). This suggests that for RNA there is either more frequent directional selection or less efficient selection against directional changes. This result is robust to the method used to classify *cis + trans* effects (Landry et al. 2005), with 63% of *cistrans* RNA features being classified as divergent for RNA vs. 22% for chromatin features (Fig, S7d: p < 2.2e-16 chi^2). Taken together, these results suggest clear differences in evolutionary trajectory between regulatory layers which reflects population processes operating at different levels of organization, as well as differences between functional gene classes.

## Discussion

We used genetic variation to better understand the impact of sequence variation in regulatory DNA on embryonic gene expression, and to shed light on how these effects are propagated or buffered through different layers of regulatory information. We generated allele-specific measurements of chromatin occupancy (ATAC-seq), chromatin activity state (ChIP-seq of chromatin modifications) and gene expression (RNA-seq) in F1 embryos from eight different genotypes across multiple stages of embryogenesis. Our analysis of this extensive dataset led to several conclusions about the impact of regulatory mutations on transcriptional phenotypes.

First, although *cis*-acting genetic variation in gene expression and associated regulatory signals is fairly common in development, its effects are not equally distributed across the genome. Allelic variation is both more frequent and has greater magnitude at distal regulatory elements (putative enhancers) compared to promoters, despite genetic variation itself being more common at promoters. This may in part be due to differences in the relative importance of sequence content at promoters and enhancers – many promoters, particularly for broadly expressed genes, are remarkably tolerant to mutations (Schor et al. 2017). Interestingly, despite having a greater magnitude, AI at distal elements is less likely to be propagated to other regulatory layers (Fig. 3), suggesting that enhancer mutations are either more effectively buffered or of lower functional impact, a hypothesis that fits well with the observed robustness of gene expression to deletions that remove distal regulatory elements (Hong et al. 2008; Cannavo et al. 2016). However, large effect gene-by-gene or gene-by-environment interactions can theoretically serve to release this ‘cryptic’ genetic variation (Gibson and Dworkin 2004; Schneider and Meyer 2017; Zheng et al. 2019). Whether such interactions are sufficiently common for regulatory traits is currently unknown, although we note here that the genetic contribution to (total count) gene expression varies considerably between time points, suggesting a substantial context-specificity to the genetic variation underlying gene expression variation.

Second, although all data types (open chromatin, histone modifications, RNA levels) are highly correlated, their explanatory values (potential causal relationships) as revealed by partial correlation analysis are far from equal. Using *cis*-acting variation as perturbations to development, we observed a strong, potentially direct relationship between genetic variants affecting open chromatin (TF binding) at both proximal and distal sites and gene expression, as expected. The relationship between histone modifications and gene expression, however, proved more surprising – in contrast to total count data, both in this study and previously reported (Lasserre et al. 2013), we note a strong, potentially causal, link between allelic-imbalance in H3K4me3 signal and allelic imbalance in associated genes.

Although highly correlated with gene expression, the functional requirement of H3K4me3 for transcription is controversial (Howe et al. 2017). Our copula analysis placed H3K4me3 downstream of RNA (Fig. 4d), suggesting that RNA levels are not impacted by variation affecting H3K4me3. This placement of RNA upstream of H3K4me3, inferred from our statistical analysis of the functional impact of genetic variation on both properties, is supported by recent genetic ablation studies showing that RNA transcription does not require H3K4me3 (Clouaire et al. 2012; Margaritis et al. 2012; Clouaire et al. 2014). This is consistent with suggestions that H3K4me3 is deposited as a consequence of transcription and may be required in more downstream post-transcriptional events (Howe et al. 2017). In addition, we also observed a second, independent, pathway in which genetic mutations affecting H3K27ac impacted gene expression, but only when they were also associated with *cis*-influenced changes in chromatin accessibility.

Third, the impact of *cis*-regulatory variation on gene expression is influenced by regulatory complexity, with genes that have more regulatory elements being less likely to show allelic imbalance (Fig 5). In part, this may be due to selection against variation in regulatory elements associated with these genes. As observed in other studies (Cannavo et al. 2016,), we found a clear reduction in allelic variation in regulatory elements associated with TFs and developmental regulators as compared to other gene categories. But selection is unlikely to be the whole story. Even when associated mutations are present, TFs and other genes with complex regulation show a degree of independence from allelic imbalance in associated regulatory layers, an active buffering process resulting from the presence of multiple regulatory inputs (Waymack 2019). Notably, the buffering of genes with multiple regulatory elements is not absolute - as the number of regulatory regions with AI near a gene increases, so does the probability that the gene shows allelic imbalance. We propose that information averaging in *cis*-regulatory landscapes enhances the overall consistency of transcriptional responses, with clustered regulatory elements, including shadow enhancers, leading to a reduction in overall allelic imbalance. In addition, large effect mutations can directly influence gene expression, with likely consequences for adaptive phenotypes, while small effect mutations, *e*.*g*. those affecting histone modifications or chromatin accessibility without affecting gene expression, may accumulate over time to have functionally relevant phenotypes.

Finally, we found that *trans*-acting variation is more common for RNA than for any other regulatory layer, with resulting consequences for the selection and heritability of gene expression relative to chromatin features. Specifically, genes with complex regulatory landscapes, *e*.*g*. transcription factors, had a higher *trans* proportion of their overall genetic influences. This observation, which is likely due to buffering effects within complex *cis* regulatory landscapes, has potentially counter intuitive evolutionary consequences, as predominantly *trans*-influenced genes are significantly more likely to show non-additive, and thus less selectable, patterns of inheritance. As a result, *trans*-acting variation affecting genes such as TFs may remain in populations even as negative selection and buffering act to reduce the influence of *cis*-acting mutations.

In summary, allelic variation in chromatin accessibility and histone modifications at regulatory elements is prevalent in the genome and capable of propagating across regulatory layers. Information flow depends on the type of regulatory element and appears mitigated at developmental factors. Notably, these *cis*-regulatory changes to individual genes do not have an appreciable effect on overall developmental programs.

## Methods

Detailed methods for all sections are provided in the supplementary file.

### Fly husbandry, crosses and embryo collection

F1 hybrid embryos were generated by crossing males from eight genetically distinct inbred lines from the *Drosophila* Genetic Reference Panel (DGRP) collection (Mackay et al. 2012) to females from a common maternal “virginizer” line. The virginizer line contains a heat-shock inducible pro-apoptotic gene (*hid*) on the Y chromosome (Starz-Gaiano et al. 2001) of a laboratory reference strain (*w*^*1118*^). We made the virginizer line isogenic by backcrossing for over 20 generations (Ghavi-Helm et al. 2019). Placing embryos from the virginizer stock at 37°C kills all male embryos, thereby facilitating the collection of a large population of isogenic virgin females, which were mated to males of different DGRP lines (listed in Fig. 1a). In addition, we collected samples from the parents of one genotype (399) for *cis*-*trans* analysis (see below).

### RNA-seq, ATAC-seq and iChIP

For three developmental stages (2-4hr, 6-8hr and 10-12hr after egg-laying), we performed RNA-Seq, ATAC-Seq, and iChIP for H3K27ac and H3K4me3 for pooled embryos of each F1 strain. All experiments were performed in biological replicates from independent embryo collections. iChIP experiments were performed as described in Lara-Astiaso et al. 2014 (Lara-Astiaso et al. 2014). ATAC-seq libraries were 125bp PE, RNA-Seq 118bp PE, and iChIP 75bp PE. In addition, gDNA from ∼100 embryos per F1 cross was extracted and 75bp SE libraries constructed. All libraries were run on a Bioanalyzer chip, multiplexed and sequenced with Illumina machines.

### Sequencing reads processing

Strain-specific genomes and liftOver chain files were constructed for each DGRP paternal line using custom scripts to insert SNPs and indels into the *Drosophila* dm3 assembly (version 5 from FlyBase). To annotate these parental genomes, we used pslMap (Zhu et al. 2007) to shift reference annotations r5.57 to the parental genomes. ATAC-seq and ChIP-seq reads were mapped using BWA (Li and Durbin 2010), while RNA-seq reads mapped using STAR (Dobin et al. 2013). In all cases, overlapping read pairs were trimmed so each base was covered only once by the higher quality read. The resulting alignments against both parental genomes were merged into a single alignment file. To generate allele-specific counts, reads were scored for their overlap with known cross-specific SNPs. Discordant reads (those overlapping alleles assigned to different parents) were discarded. Genomic DNA was generated for each F1 line to filter potentially miscalled variants, and simulated reads from each parental genome were used to assess and filter out regions with likely mappability errors. Peak calling was performed using Macs2 (-broad) for iChIP reads and Hotspot for ATAC-seq reads (Zhang et al. 2008; John et al. 2011). To compare between lines and times, we constructed merged peak coordinates across samples (Supplementary methods).

### Test for allele-specific imbalance

Due to the extensive maternally deposited transcripts still present at 2-4 hours, we excluded the RNA-seq data from this time point from all down-stream allele-specific analysis to avoid potential confounding effects in allelic imbalance measurements. To test for allelic imbalance, an empirical Bayesian method was used to test the null hypothesis for differences in read counts between F1 alleles for each feature of each data set (RNA-seq, ATAC-seq, H3K4me3, H3K27ac). Individual tests were performed for each line and for each time point. Total F1 counts 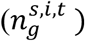 can be modeled on an allele-specific basis 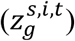 using a beta-binomial distribution. Specifically, 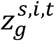 denotes the number of reads from the maternal allele mapped to feature f for pool of individuals i, of paternal strain s, at time t. 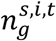 denotes the total number of reads mapping to genes for pool of individuals i of strain s, at time t.

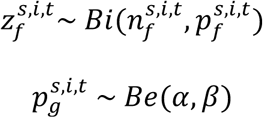

where *α, β* are the shape parameters of the beta distribution. We tested the following scenarios by maximum likelihood estimation:

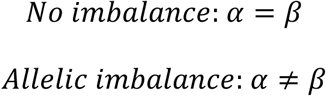

Due to limited replicates per condition, we used information across features to reduce the uncertainty of estimates and improve power by assuming that all features have the same mean-variance relationship (Robinson et al. 2010; Love et al. 2014). Empirical data was used to estimate the over-dispersion parameter (*ρ*) for each data type based on the beta-binomial distribution. Maximum likelihood estimation was used to obtain *α* and *β* for each feature of time *t* and strain *s. ρ* is calculated as follows:

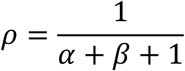

The mean over-dispersion value for all features was used as the shrinkage term and likelihood ratio tests (df=1) were used to obtain a p-value, which was adjusted using FDR (Benjamini 1995). Autosomes were tested separately to sex chromosomes; features on chromosome X were tested using a background allelic ratio of no imbalance centered on the averaged ratio of maternal versus paternal alleles across the data set being compared (i.e. RNA, ATAC, H3K4me3, H3K27ac). Autosomal features were tested using a null distribution of 0.5.

### Allele-specific changes across lines and developmental time

A linear mixed-effects model, where random effect components were incorporated, was used to estimate variability between pools of individuals, time points and lines,

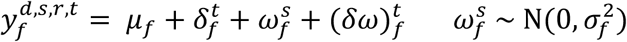

*μ*_*f*_ is the intercept term. 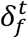 is a random effect term denoting time. 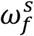 is a random effect based on strain and 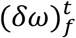 is a interaction term for time by strain.

To infer the significance of time or strain dependent allele bias, we restricted the values that the parameters can take. Library size differences were corrected for at the allele-combined count level using the TMM method in ‘edgeR’ (Robinson et al. 2010) prior to analysis. Count data was filtered for reads with more than 20 allele-combined counts. Each autosomal feature was tested using read counts at SNPs common to all lines. Not all features contained enough information for statistical testing, subsequent analyses were limited to features with at least six samples in each of the three time points in at least four genetic strain.

### Allele-specific changes across regulatory layers

Intersection-union tests were used to examine the pairwise co-occurrence of allelic imbalance in overlapping genes/features, limited to autosomes, based on rejecting the null hypothesis if a significant outcome with respect to the feature compared at the same time point exists for both data types (Berger 1996).

To infer pairwise relationships between regulatory data types while reducing indirect relations, partial correlation analysis was performed using ‘GeneNet’ (Opgen-Rhein and Strimmer 2007) for both allelic ratios and total count data. Directional dependence modeling was performed in a regression framework using copulas to describe the bivariate distribution between our pairwise datasets (Lee and Kim 2019). Copula regression was used to infer the flow of information for pairwise relationships that showed a significant relationship in partial correlation analyses.

Conditional probabilities for the probability of allelic imbalance given imbalance in a different regulatory data type were calculated by the definition:

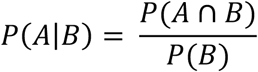

where *A* and *B* are the probabilities of allelic imbalance in each data type.

### *Ci*s *trans* analysis

For one F1 line (vgn x 399) and its parental lines, maximum likelihood estimation (MLE) was used to compare parental and offspring ratios simultaneously to determine whether gene expression, chromatin accessibility, H3K4me3 and H3K27ac enrichments are influenced by *cis*-, *trans*-, conserved or both *cis*- and *trans*-acting effects by modeling read counts. For parents, the data was modeled using negative binomial distributions and allelic differences in F1 alleles modeled using beta-binomial distribution (Supplemental Methods). We constrained parameter estimation for each model based on four different regulatory scenarios and derived maximum likelihood values for each hypothetical case on a site-by-site basis. In the presentation of the proportions of features assigned to each category (Fig. 6a, S7c), we presented the maximum likelihood assignment. In subsequent analyses, we limited analyses to features that showed a BIC difference >= 2.

### Test for compensatory mutation

Genes were classified as having *cis*- and *trans*-acting influences following the procedure of Goncalves et al. (Goncalves et al. 2012). For all genes, we asked if their *cis* and *trans* contributes act to reinforce one another (same direction) or if they operated in opposite directions. Formally, for the i_th gene, we define the average log2 fold change for the parental lines as x_i and the average log2 allelic ratio from the F1 data as y_i. We then classified:

Opposite – cis stronger: (0 < y_i < x_i) OR (0 > y_i > x_i)

Opposite – trans stronger: (x_i < 0 < y_i) OR (y_i < 0 < x_i)

Same – cis stronger (0 < x_i < y_i < 2x_i) OR (0 > x_i > y_i > 2x_i)

Same – trans stronger (0 < 2x_i < y_i) OR (0 > 2x_i > y_i)

A complementary analysis following Landry et al (Landry et al. 2005) can be found in the supplemental methods.

### Measuring additive vs. non-additive heritability

In the case of additively inherited gene expression (or read counts for any of our measured features), the signal observed in the F1 is expected to be equal to the midpoint (average) of the two parents, while non-additively inherited genes/features should show a significant departure from that midpoint. To formally test for non-additivity, we made use of the standard workflow in DESeq2 with two modifications. First, we set the betaPrior option equal to TRUE. After setting the reference genotype to the F1 (vgn x 399) using the relevel function, we then extracted the results using the ‘results’ function and the contrast vector c(0,1,-.5, -.5) to contrast the full value of the F1 genotype with ½(vgn + 399). Features with an FDR < .1 were considered as “non-additive”.

## Data Access

All raw data has been deposited to EMBL-EBI hosted ArrayExpress, accession numbers: E-MTAB-8877 (gDNA), E-MTAB-8878 (RNA-seq), E-MTAB-8879 (ATAC-seq), E-MTAB-8880 (ChIP-seq H3K4me3, H3K27ac). Processed data, including total counts, allelic ratios, cis/trans estimates, estimated per-feature heritability, mappability filters, and parental genotype files can all be downloaded from http://furlonglab.embl.de/data

## Acknowledgements

We are very grateful to members of the Furlong lab for discussions and comments, in particular to Olga Sigalova, Adam Rabinowitz, Marijn van Jaarsveld and Matteo Perino. This work was technically supported by the EMBL Genomics Core Facility and the public resources of FlyBase, BDGP, and RedFly. The work was financially supported by the European Research Council (ERC advanced grant) agreement 322851 (CisRegVar) and 787611 (DeCRyPT) to E.E.F.

## Author Contributions

EF, DG, and BZ conceived the project. BZ developed the ATAC-seq and RRV the iChIP protocol for *Drosophila* embryos. BZ and RRV generated the data with help from DG. DG, SF, and EW performed data analysis. DG and SF performed mapping bias analysis and allelic and total count data processing. SF performed partial correlation analysis, and EW the statistical modeling for allelic imbalance and Copula analysis. DG and EW performed cis/trans analysis. DG performed analysis of heritability and evolutionary variation. EF, DG, EW, BZ and SF wrote the manuscript with input from all authors. MT and DF helped to revise the manuscript.

## References

Ahituv N, Zhu Y, Visel A, Holt A, Afzal V, Pennacchio LA, Rubin EM. 2007. Deletion of ultraconserved elements yields viable mice. PLoS Biol 5: e234.

Battlay P, Schmidt JM, Fournier-Level A, Robin C. 2016. Genomic and Transcriptomic Associations Identify a New Insecticide Resistance Phenotype for the Selective Sweep at the Cyp6g1 Locus of Drosophila melanogaster. G3 (Bethesda) 6: 2573–2581.

Battle A, Khan Z, Wang SH, Mitrano A, Ford MJ, Pritchard JK, Gilad Y. 2015. Genomic variation. Impact of regulatory variation from RNA to protein. Science 347: 664–667.

Behera V, Evans P, Face CJ, Hamagami N, Sankaranarayanan L, Keller CA, Giardine B, Tan K, Hardison RC, Shi J et al. 2018. Exploiting genetic variation to uncover rules of transcription factor binding and chromatin accessibility. Nat Commun 9: 782.

Benjamini Y, and Hochberg, Y. 1995. Controlling the False Discovery Rate: A Practical and Powerful Approach to Multiple Testing. Journal of the Royal Statistical Society: Series B (Methodological) 57: 289–300.

Berger RLaH, Jason C. 1996. Bioequivalence trials, intersection-union tests and equivalence confidence sets. Statistical Science 11: 283–302.

Bonn S, Zinzen RP, Girardot C, Gustafson EH, Perez-Gonzalez A, Delhomme N, Ghavi-Helm Y, Wilczynski B, Riddell A, Furlong EE. 2012. Tissue-specific analysis of chromatin state identifies temporal signatures of enhancer activity during embryonic development. Nat Genet 44: 148–156.

Borok MJ, Tran DA, Ho MC, Drewell RA. 2010. Dissecting the regulatory switches of development: lessons from enhancer evolution in Drosophila. Development 137: 5–13.

Brown CD, Johnson DS, Sidow A. 2007. Functional architecture and evolution of transcriptional elements that drive gene coexpression. Science 317: 1557–1560.

Buenrostro JD, Giresi PG, Zaba LC, Chang HY, Greenleaf WJ. 2013. Transposition of native chromatin for fast and sensitive epigenomic profiling of open chromatin, DNA-binding proteins and nucleosome position. Nat Methods 10: 1213–1218.

Bullaughey K. 2011. Changes in selective effects over time facilitate turnover of enhancer sequences. Genetics 187: 567–582.

Cannavo E, Khoueiry P, Garfield DA, Geeleher P, Zichner T, Gustafson EH, Ciglar L, Korbel JO, Furlong EE. 2016. Shadow Enhancers Are Pervasive Features of Developmental Regulatory Networks. Curr Biol 26: 38–51.

Cannavo E, Koelling N, Harnett D, Garfield D, Casale FP, Ciglar L, Gustafson HE, Viales RR, Marco-Ferreres R, Degner JF et al. 2017. Genetic variants regulating expression levels and isoform diversity during embryogenesis. Nature 541: 402–406.

Celniker SE, Dillon LA, Gerstein MB, Gunsalus KC, Henikoff S, Karpen GH, Kellis M, Lai EC, Lieb JD, MacAlpine DM et al. 2009. Unlocking the secrets of the genome. Nature 459: 927–930.

Chen L, Ge B, Casale FP, Vasquez L, Kwan T, Garrido-Martin D, Watt S, Yan Y, Kundu K, Ecker S et al. 2016. Genetic Drivers of Epigenetic and Transcriptional Variation in Human Immune Cells. Cell 167: 1398–1414 e1324.

Clouaire T, Webb S, Bird A. 2014. Cfp1 is required for gene expression-dependent H3K4 trimethylation and H3K9 acetylation in embryonic stem cells. Genome Biol 15: 451.

Clouaire T, Webb S, Skene P, Illingworth R, Kerr A, Andrews R, Lee JH, Skalnik D, Bird A. 2012. Cfp1 integrates both CpG content and gene activity for accurate H3K4me3 deposition in embryonic stem cells. Genes Dev 26: 1714–1728.

Conrad T, Cavalli FM, Vaquerizas JM, Luscombe NM, Akhtar A. 2012. Drosophila dosage compensation involves enhanced Pol II recruitment to male X-linked promoters. Science 337: 742–746.

Core LJ, Waterfall JJ, Gilchrist DA, Fargo DC, Kwak H, Adelman K, Lis JT. 2012. Defining the status of RNA polymerase at promoters. Cell Rep 2: 1025–1035.

Cretekos CJ, Wang Y, Green ED, Martin JF, Rasweiler JJt, Behringer RR. 2008. Regulatory divergence modifies limb length between mammals. Genes Dev 22: 141–151.

Cusanovich DA, Reddington JP, Garfield DA, Daza RM, Aghamirzaie D, Marco-Ferreres R, Pliner HA, Christiansen L, Qiu X, Steemers FJ et al. 2018. The cis-regulatory dynamics of embryonic development at single-cell resolution. Nature 555: 538–542.

Daborn P, Boundy S, Yen J, Pittendrigh B, ffrench-Constant R. 2001. DDT resistance in Drosophila correlates with Cyp6g1 over-expression and confers cross-resistance to the neonicotinoid imidacloprid. Mol Genet Genomics 266: 556–563.

Dobin A, Davis CA, Schlesinger F, Drenkow J, Zaleski C, Jha S, Batut P, Chaisson M, Gingeras TR. 2013. STAR: ultrafast universal RNA-seq aligner. Bioinformatics 29: 15–21.

Doitsidou M, Flames N, Topalidou I, Abe N, Felton T, Remesal L, Popovitchenko T, Mann R, Chalfie M, Hobert O. 2013. A combinatorial regulatory signature controls terminal differentiation of the dopaminergic nervous system in C. elegans. Genes Dev 27: 1391–1405.

Epstein DJ. 2009. Cis-regulatory mutations in human disease. Brief Funct Genomic Proteomic 8: 310–316.

Frankel N, Davis GK, Vargas D, Wang S, Payre F, Stern DL. 2010. Phenotypic robustness conferred by apparently redundant transcriptional enhancers. Nature 466: 490–493.

Garfield DA, Runcie DE, Babbitt CC, Haygood R, Nielsen WJ, Wray GA. 2013. The impact of gene expression variation on the robustness and evolvability of a developmental gene regulatory network. PLoS Biol 11: e1001696.

Georgiev P, Chlamydas S, Akhtar A. 2011. Drosophila dosage compensation: males are from Mars, females are from Venus. Fly (Austin) 5: 147–154.

Ghavi-Helm Y, Jankowski A, Meiers S, Viales RR, Korbel JO, Furlong EEM. 2019. Highly rearranged chromosomes reveal uncoupling between genome topology and gene expression. Nat Genet 51: 1272–1282.

Gibson G, Dworkin I. 2004. Uncovering cryptic genetic variation. Nat Rev Genet 5: 681–690.

Goncalves A, Leigh-Brown S, Thybert D, Stefflova K, Turro E, Flicek P, Brazma A, Odom DT, Marioni JC. 2012. Extensive compensatory cis-trans regulation in the evolution of mouse gene expression. Genome Res 22: 2376–2384.

Hong JW, Hendrix DA, Levine MS. 2008. Shadow enhancers as a source of evolutionary novelty. Science 321: 1314.

Howe FS, Fischl H, Murray SC, Mellor J. 2017. Is H3K4me3 instructive for transcription activation? Bioessays 39: 1–12.

John S, Sabo PJ, Thurman RE, Sung MH, Biddie SC, Johnson TA, Hager GL, Stamatoyannopoulos JA. 2011. Chromatin accessibility pre-determines glucocorticoid receptor binding patterns. Nat Genet 43: 264–268.

Junion G, Spivakov M, Girardot C, Braun M, Gustafson EH, Birney E, Furlong EE. 2012. A transcription factor collective defines cardiac cell fate and reflects lineage history. Cell 148: 473–486.

Karlic R, Chung HR, Lasserre J, Vlahovicek K, Vingron M. 2010. Histone modification levels are predictive for gene expression. Proc Natl Acad Sci U S A 107: 2926–2931.

Kasowski M, Grubert F, Heffelfinger C, Hariharan M, Asabere A, Waszak SM, Habegger L, Rozowsky J, Shi M, Urban AE et al. 2010. Variation in transcription factor binding among humans. Science 328: 232–235.

Kheradpour P, Ernst J, Melnikov A, Rogov P, Wang L, Zhang X, Alston J, Mikkelsen TS, Kellis M. 2013. Systematic dissection of regulatory motifs in 2000 predicted human enhancers using a massively parallel reporter assay. Genome Res 23: 800–811.

Khoueiry P, Girardot C, Ciglar L, Peng PC, Gustafson EH, Sinha S, Furlong EE. 2017. Uncoupling evolutionary changes in DNA sequence, transcription factor occupancy and enhancer activity. Elife 6: e28440.

Kilpinen H, Waszak SM, Gschwind AR, Raghav SK, Witwicki RM, Orioli A, Migliavacca E, Wiederkehr M, Gutierrez-Arcelus M, Panousis NI et al. 2013. Coordinated effects of sequence variation on DNA binding, chromatin structure, and transcription. Science 342: 744–747.

Kim JM, Jung YS, Sungur EA, Han KH, Park C, Sohn I. 2008. A copula method for modeling directional dependence of genes. BMC Bioinformatics 9: 225.

Kircher M, Xiong C, Martin B, Schubach M, Inoue F, Bell RJA, Costello JF, Shendure J, Ahituv N. 2019. Saturation mutagenesis of twenty disease-associated regulatory elements at single base-pair resolution. Nat Commun 10: 3583.

Knowles DA, Davis JR, Edgington H, Raj A, Fave MJ, Zhu X, Potash JB, Weissman MM, Shi J, Levinson DF et al. 2017. Allele-specific expression reveals interactions between genetic variation and environment. Nat Methods 14: 699–702.

Kvon EZ, Kazmar T, Stampfel G, Yanez-Cuna JO, Pagani M, Schernhuber K, Dickson BJ, Stark A. 2014. Genome-scale functional characterization of Drosophila developmental enhancers in vivo. Nature 512: 91–95.

Kwasnieski JC, Fiore C, Chaudhari HG, Cohen BA. 2014. High-throughput functional testing of ENCODE segmentation predictions. Genome Res 24: 1595–1602.

Landry CR, Wittkopp PJ, Taubes CH, Ranz JM, Clark AG, Hartl DL. 2005. Compensatory cis-trans evolution and the dysregulation of gene expression in interspecific hybrids of Drosophila. Genetics 171: 1813–1822.

Lara-Astiaso D, Weiner A, Lorenzo-Vivas E, Zaretsky I, Jaitin DA, David E, Keren-Shaul H, Mildner A, Winter D, Jung S et al. 2014. Immunogenetics. Chromatin state dynamics during blood formation. Science 345: 943–949.

Lasserre J, Chung HR, Vingron M. 2013. Finding associations among histone modifications using sparse partial correlation networks. PLoS Comput Biol 9: e1003168.

Lee N, Kim JM. 2019. Copula directional dependence for inference and statistical analysis of whole-brain connectivity from fMRI data. Brain Behav 9: e01191.

Lemos B, Araripe LO, Fontanillas P, Hartl DL. 2008. Dominance and the evolutionary accumulation of cis- and trans-effects on gene expression. Proc Natl Acad Sci U S A 105: 14471–14476.

Li H, Durbin R. 2010. Fast and accurate long-read alignment with Burrows-Wheeler transform. Bioinformatics 26: 589–595.

Long HK, Prescott SL, Wysocka J. 2016. Ever-Changing Landscapes: Transcriptional Enhancers in Development and Evolution. Cell 167: 1170–1187.

Love MI, Huber W, Anders S. 2014. Moderated estimation of fold change and dispersion for RNA-seq data with DESeq2. Genome Biol 15: 550.

Lowe WL, Jr., Reddy TE. 2015. Genomic approaches for understanding the genetics of complex disease. Genome Res 25: 1432–1441.

Lu R, Rogan PK. 2018. Transcription factor binding site clusters identify target genes with similar tissue-wide expression and buffer against mutations. F1000Res 7: 1933.

Lucchesi JC, Kuroda MI. 2015. Dosage compensation in Drosophila. Cold Spring Harb Perspect Biol 7: a019398.

Lynch M, Walsh B. 1998. Genetics and analysis of quantitative traits. Sinauer, Sunderland, Mass.

Mackay TF, Richards S, Stone EA, Barbadilla A, Ayroles JF, Zhu D, Casillas S, Han Y, Magwire MM, Cridland JM et al. 2012. The Drosophila melanogaster Genetic Reference Panel. Nature 482: 173–178.

Margaritis T, Oreal V, Brabers N, Maestroni L, Vitaliano-Prunier A, Benschop JJ, van Hooff S, van Leenen D, Dargemont C, Geli V et al. 2012. Two distinct repressive mechanisms for histone 3 lysine 4 methylation through promoting 3’-end antisense transcription. PLoS Genet 8: e1002952.

Meiklejohn CD, Coolon JD, Hartl DL, Wittkopp PJ. 2014. The roles of cis- and trans-regulation in the evolution of regulatory incompatibilities and sexually dimorphic gene expression. Genome Res 24: 84–95.

Mi H, Vandergriff J, Campbell M, Narechania A, Majoros W, Lewis S, Thomas PD, Ashburner M. 2003. Assessment of genome-wide protein function classification for Drosophila melanogaster. Genome Res 13: 2118–2128.

Mikhaylichenko O, Bondarenko V, Harnett D, Schor IE, Males M, Viales RR, Furlong EEM. 2018. The degree of enhancer or promoter activity is reflected by the levels and directionality of eRNA transcription. Genes Dev 32: 42–57.

Montavon T, Soshnikova N, Mascrez B, Joye E, Thevenet L, Splinter E, de Laat W, Spitz F, Duboule D. 2011. A regulatory archipelago controls Hox genes transcription in digits. Cell 147: 1132–1145.

Moyerbrailean GA, Richards AL, Kurtz D, Kalita CA, Davis GO, Harvey CT, Alazizi A, Watza D, Sorokin Y, Hauff N et al. 2016. High-throughput allele-specific expression across 250 environmental conditions. Genome Res 26: 1627–1638.

Opgen-Rhein R, Strimmer K. 2007. From correlation to causation networks: a simple approximate learning algorithm and its application to high-dimensional plant gene expression data. BMC Syst Biol 1: 37.

Paaby AB, Gibson G. 2016. Cryptic Genetic Variation in Evolutionary Developmental Genetics. Biology (Basel) 5: E28.

Pai AA, Pritchard JK, Gilad Y. 2015. The genetic and mechanistic basis for variation in gene regulation. PLoS Genet 11: e1004857.

Pal K, Forcato M, Jost D, Sexton T, Vaillant C, Salviato E, Mazza EMC, Lugli E, Cavalli G, Ferrari F. 2019. Global chromatin conformation differences in the Drosophila dosage compensated chromosome X. Nat Commun 10: 5355.

Pradeepa MM, Grimes GR, Kumar Y, Olley G, Taylor GC, Schneider R, Bickmore WA. 2016. Histone H3 globular domain acetylation identifies a new class of enhancers. Nat Genet 48: 681–686.

Reddy TE, Gertz J, Pauli F, Kucera KS, Varley KE, Newberry KM, Marinov GK, Mortazavi A, Williams BA, Song L et al. 2012. Effects of sequence variation on differential allelic transcription factor occupancy and gene expression. Genome Res 22: 860–869.

Robinson MD, McCarthy DJ, Smyth GK. 2010. edgeR: a Bioconductor package for differential expression analysis of digital gene expression data. Bioinformatics 26: 139–140.

Schneider RF, Meyer A. 2017. How plasticity, genetic assimilation and cryptic genetic variation may contribute to adaptive radiations. Mol Ecol 26: 330–350.

Schor IE, Degner JF, Harnett D, Cannavo E, Casale FP, Shim H, Garfield DA, Birney E, Stephens M, Stegle O et al. 2017. Promoter shape varies across populations and affects promoter evolution and expression noise. Nat Genet 49: 550–558.

Spitz F, Furlong EE. 2012. Transcription factors: from enhancer binding to developmental control. Nat Rev Genet 13: 613–626.

Spivakov M, Akhtar J, Kheradpour P, Beal K, Girardot C, Koscielny G, Herrero J, Kellis M, Furlong EE, Birney E. 2012. Analysis of variation at transcription factor binding sites in Drosophila and humans. Genome Biol 13: R49.

Starz-Gaiano M, Cho NK, Forbes A, Lehmann R. 2001. Spatially restricted activity of a Drosophila lipid phosphatase guides migrating germ cells. Development 128: 983–991.

Tirosh I, Reikhav S, Levy AA, Barkai N. 2009. A yeast hybrid provides insight into the evolution of gene expression regulation. Science 324: 659–662.

Turner LM, Chuong EB, Hoekstra HE. 2008. Comparative analysis of testis protein evolution in rodents. Genetics 179: 2075–2089.

Uhl JD, Zandvakili A, Gebelein B. 2016. A Hox Transcription Factor Collective Binds a Highly Conserved Distal-less cis-Regulatory Module to Generate Robust Transcriptional Outcomes. PLoS Genet 12: e1005981.

Urban J, Kuzu G, Bowman S, Scruggs B, Henriques T, Kingston R, Adelman K, Tolstorukov M, Larschan E. 2017. Enhanced chromatin accessibility of the dosage compensated Drosophila male X-chromosome requires the CLAMP zinc finger protein. PLoS One 12: e0186855.

Villar D, Berthelot C, Aldridge S, Rayner TF, Lukk M, Pignatelli M, Park TJ, Deaville R, Erichsen JT, Jasinska AJ et al. 2015. Enhancer evolution across 20 mammalian species. Cell 160: 554–566.

Waszak SM, Delaneau O, Gschwind AR, Kilpinen H, Raghav SK, Witwicki RM, Orioli A, Wiederkehr M, Panousis NI, Yurovsky A et al. 2015. Population Variation and Genetic Control of Modular Chromatin Architecture in Humans. Cell 162: 1039–1050.

Waymack RF, A.; Enciso, G.; Wunderlich, Z.. 2019. Shadow enhancers suppress input transcription factor noise through distinct regulatory logic. bioRxiv doi: 10.1101/778092.

Wittkopp PJ, Haerum BK, Clark AG. 2004. Evolutionary changes in cis and trans gene regulation. Nature 430: 85–88.

Wittkopp PJ, Kalay G. 2011. Cis-regulatory elements: molecular mechanisms and evolutionary processes underlying divergence. Nat Rev Genet 13: 59–69.

Wong ES, Schmitt BM, Kazachenka A, Thybert D, Redmond A, Connor F, Rayner TF, Feig C, Ferguson-Smith AC, Marioni JC et al. 2017. Interplay of cis and trans mechanisms driving transcription factor binding and gene expression evolution. Nat Commun 8: 1092.

Xiong N, Kang C, Raulet DH. 2002. Redundant and unique roles of two enhancer elements in the TCRgamma locus in gene regulation and gammadelta T cell development. Immunity 16: 453–463.

Yadav A, Dhole K, Sinha H. 2016. Differential Regulation of Cryptic Genetic Variation Shapes the Genetic Interactome Underlying Complex Traits. Genome Biol Evol 8: 3559–3573.

Zabidi MA, Arnold CD, Schernhuber K, Pagani M, Rath M, Frank O, Stark A. 2015. Enhancer-core-promoter specificity separates developmental and housekeeping gene regulation. Nature 518: 556–559.

Zhang Y, Liu T, Meyer CA, Eeckhoute J, Johnson DS, Bernstein BE, Nusbaum C, Myers RM, Brown M, Li W et al. 2008. Model-based analysis of ChIP-Seq (MACS). Genome Biol 9: R137.

Zheng J, Payne JL, Wagner A. 2019. Cryptic genetic variation accelerates evolution by opening access to diverse adaptive peaks. Science 365: 347–353.

Zheng W, Zhao H, Mancera E, Steinmetz LM, Snyder M. 2010. Genetic analysis of variation in transcription factor binding in yeast. Nature 464: 1187–1191.

Zhu J, Sanborn JZ, Diekhans M, Lowe CB, Pringle TH, Haussler D. 2007. Comparative genomics search for losses of long-established genes on the human lineage. PLoS Comput Biol 3: e247.

Zinzen RP, Girardot C, Gagneur J, Braun M, Furlong EE. 2009. Combinatorial binding predicts spatio-temporal cis-regulatory activity. Nature 462: 65–70.

